# Presence of endogenous viral elements negatively correlates with FeLV susceptibility in puma and domestic cat cells

**DOI:** 10.1101/2020.06.23.168351

**Authors:** Elliott S. Chiu, Sue VandeWoude

## Abstract

While feline leukemia virus (FeLV) has been shown to infect felid species other than the endemic domestic cat host, differences in FeLV susceptibility among species has not been evaluated. Previous reports have noted a negative correlation between enFeLV copy number and exogenous FeLV infection outcomes in domestic cats. Since felids outside the genus *Felis* do not harbor enFeLV genomes, we hypothesized absence of enFeLV results in more severe disease consequences in felid species lacking these genomic elements. We infected primary fibroblasts isolated from domestic cats (*Felis catus*) and pumas (*Puma concolor*) with FeLV and quantitated proviral and viral antigen loads. Domestic cat enFeLV *env* and LTR copy numbers were determined for each individual and compared to FeLV viral outcomes. FeLV proviral and antigen levels were also measured in 6 naturally infected domestic cats and 11 naturally infected Florida panthers (*P. concolor coryi*). We demonstrated that puma fibroblasts are more permissive to FeLV than domestic cat cells, and domestic cat FeLV restriction was highly related to enFeLV LTR copy number. Terminal tissues from FeLV-infected Florida panthers and domestic cats had similar exFeLV proviral copy numbers, but Florida panther tissues have higher FeLV antigen loads. Our work indicates enFeLV LTR elements negatively regulate exogenous FeLV replication. Further, *Puma concolor* lacking enFeLV are more permissive to FeLV infection than domestic cats, suggesting endogenization can play a beneficial role in mitigating exogenous retroviral infections. Conversely, presence of endogenous retroelements may relate to new host susceptibility during viral spillover events.

**Importance:** Feline leukemia virus (FeLV) can infect a variety of felid species. Only the primary domestic cat host and related small cat species harbor a related endogenous virus in their genomes. Previous studies noted a negative association between the endogenous virus copy number and exogenous virus infection in domestic cats. This report shows that puma cells, which lack endogenous FeLV, produce more virus more rapidly than domestic cat fibroblasts following cell culture challenge. We document a strong association between domestic cat cell susceptibility and FeLV long terminal repeat (LTR) copy number, similar to observations in natural FeLV infections. Viral replication does not, however, correlate with FeLV *env* copy number, suggesting this effect is specific to FeLV LTR elements. This discovery indicates a protective capacity of the endogenous virus against the exogenous form, either via direct interference or indirectly via gene regulation, and may suggest evolutionary outcomes of retroviral endogenization.

## Introduction

The vast majority of vertebrate genomes, including up to 8% of the human genome, harbor fossils of ancient viral infections made up predominantly of retroviral genetic material (1-3). During infection, the retroviral RNA genome is reverse-transcribed to form double-stranded DNA, which is in turn integrated into the host’s genome (4). While most of these infections target somatic cells, these viruses are capable of infecting and integrating into germ cells (5). The consequences of viral integration into the germline is profound, and ultimately the virus is vertically transmitted as permanent genetic elements inherited in a Mendelian fashion (6). Fixation of the retroviral content in host genomes is a process termed endogenization and leads to new host genetic elements called endogenous retroviruses (ERVs) (7). ERVs in their early stages are believed to undergo massive changes during host cell transcription, and the foreign, potentially deleterious genetic material accumulates mutations and deletions that often render the newly endogenized virus defunct (8). While typically unable to produce infectious virions, many ERVs are still capable of undergoing transcription and may even produce functional viral proteins. Certain ERVs are known to function in important physiologic, cellular, or biological processes, including placentation, oncogenesis, immune modulation, and infectious disease progression (9, 10).

Endogenous feline leukemia virus (enFeLV) is an example of an ERV which has a horizontally transmitted retroviral counterpart (feline leukemia virus, FeLV). Only members of the *Felis* genus harbor enFeLV as endogenization is believed to have originated after the *Felis* genus split off from other members of the Felidae family (11, 12). FeLV can infect felid species that harbor enFeLV (i.e., domestic cats) as well as species that lack enFeLV (i.e., puma). FeLV epizootics have been documented in multiple non-*Felis* species populations including the North American puma (*Puma concolor*) (13-19).

FeLV represents an endogenous-exogenous retroviral system that has perhaps been best studied with regard to disease biology and outcome during naturally occurring infections in an outbred, highly dispersed mammalian host. Thus, evaluation of this system provides opportunities to better understand ERV-exogenous viral interactions that are highly relevant to virus and host evolution and ecology. FeLV epizootics in wild felids are characterized by serious disease of epidemic proportions, (20, 21), whereas FeLV infection in adult cats frequently results in regressive and abortive infections (22). It has been hypothesized that enFeLV may be associated with differences in infection outcome. We previously demonstrated enFeLV long terminal repeat (LTR) copy number was associated with better infection outcomes during a natural FeLV outbreak in a multi-cat household (23). Here, we evaluate FeLV infection of puma (*P. concolor*) and domestic cat cells *in vitro* and *in situ* to examine the susceptibility of endemic and novel hosts to FeLV infection with respect to enFeLV to provide further evaluation of this relationship.

## Results

Primary fibroblasts were successfully propagated from ear punches from three free-ranging puma (two kittens of unknown sex and one adult male) and 1 abdominal skin incision from a mature adult female captive puma. Primary fibroblasts were cultured from 7 domestic cats abdominal full skin biopsies from necropsied cats at Colorado State University (6 male, 1 female).

### Fibroblast infections

Exogenous FeLV proviral load was significantly and substantially greater in puma fibroblasts compared to domestic cat cells (Figure 1A). At days 5 and 10, puma cells were infected with approximately 1 FeLV provirus per cell, whereas domestic cat cell infections were approximately a log lower. Experimental control FeLV infections of Crandall Reeves feline kidney cells (CrFK) displayed reproducible levels of infections with tight interquartile ranges while domestic cat and puma fibroblast infections displayed greater individual variance (Figure 1A). FeLV proviral load was significantly higher in puma versus domestic cat cells (Kruskal-Wallis test; *Fca* D5 vs. *Pco* D5 H = −29.8, p<0.01; *Fca* D10 vs. *Pco* D10 H = −32.55, p<0.01). FeLV viral antigen production was similarly greater in puma fibroblasts (Figure 1B). Puma cell-generated FeLV antigen detected in the supernatant was greater than in domestic cat cultures beginning days 3 and continued to be greater for the rest of the experiment (Repeated measures ANOVA; F=8.78 p=0.001).

**Figure 1.**
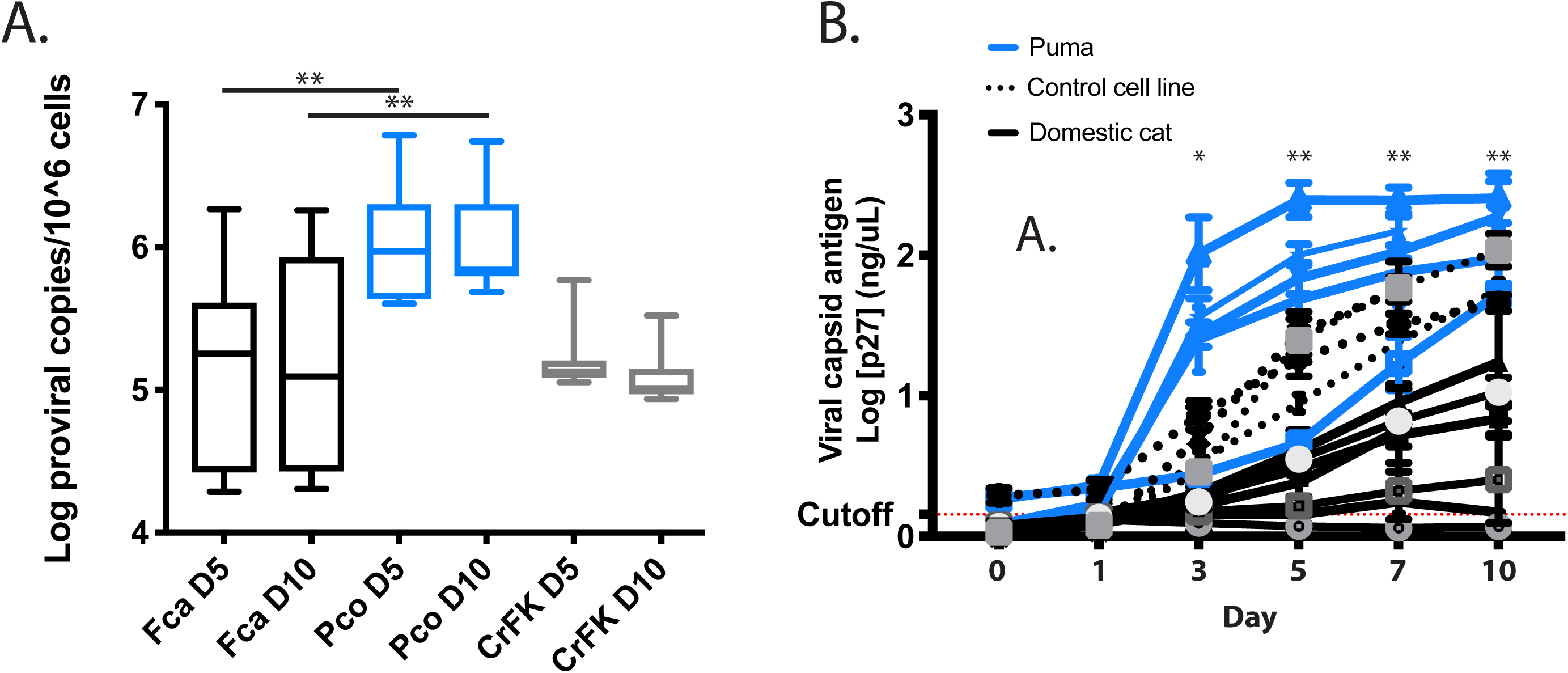
**A) Puma fibroblasts exposed to FeLV have greater FeLV proviral load compared to domestic cat fibroblasts.** On both day 5 and day 10, puma cells (Pco) demonstrated significantly higher proviral copy numbers than domestic cat primary fibroblasts (Kruskal-Wallis; **=p<0.01). Variance between domestic cats (Fca) cells was greater than that of puma cells, while CrFK cell infections displayed the least amount of variation. **B) FeLV viral capsid antigen (p27) production is higher in puma versus domestic cat cells.** Beginning day 3 PI, puma cells replicated more virus than domestic cat cells (Repeated measure ANOVA; individual test statistics *=p<0.05, **=p<0.01). One puma, one domestic cat, and one control cat experiment were ended at day 7. Cutoff values were established by 3x standard error above average value for negative control wells (red line).

Domestic cat and puma viable cell counts were equivalent on days 5 or 10, averaging 1.28×10^5^ cells per 2 cm^2^ for domestic cat cells and 1.92×10^5^ cells per 2 cm^2^ for puma cells (Supplemental figure 1A). CrFKs are smaller than domestic cat and puma primary fibroblasts and therefore cell density was higher in these cultures (3.54 ×10^5^ cells per 2 cm^2^ at day 5 and 4.13 ×10^5^ cells per 2 cm^2^ at day 10). Percent dead cells (as measured by trypan blue exclusion) on days 5 and 10 ranged between 3.94±0.70% in for domestic cat cells and 4.73%±1.45% for puma cells regardless of infection status (Supplemental figure 1B). There was a consistent trend for lower percent mortality at day 5 when cells first reached confluency. CrFKs experienced greater cell mortality over the course of the infection regardless of infection status (2.59% at day 5 and 14.26% at day 10).

### enFeLV-exFeLV correlation

As anticipated, domestic cat cells harbored more enFeLV LTRs than enFeLV *env* genes (Figure 2A). Normalized LTR sequence copy numbers in domestic cat cells ranged from 32 and 74 copies per cell with an average of 57 copies per cell. Copy numbers for enFeLV *env* were significantly lower, ranging from 9 to 13 copies per cell with an average of 11 copies.

**Figure 2.**
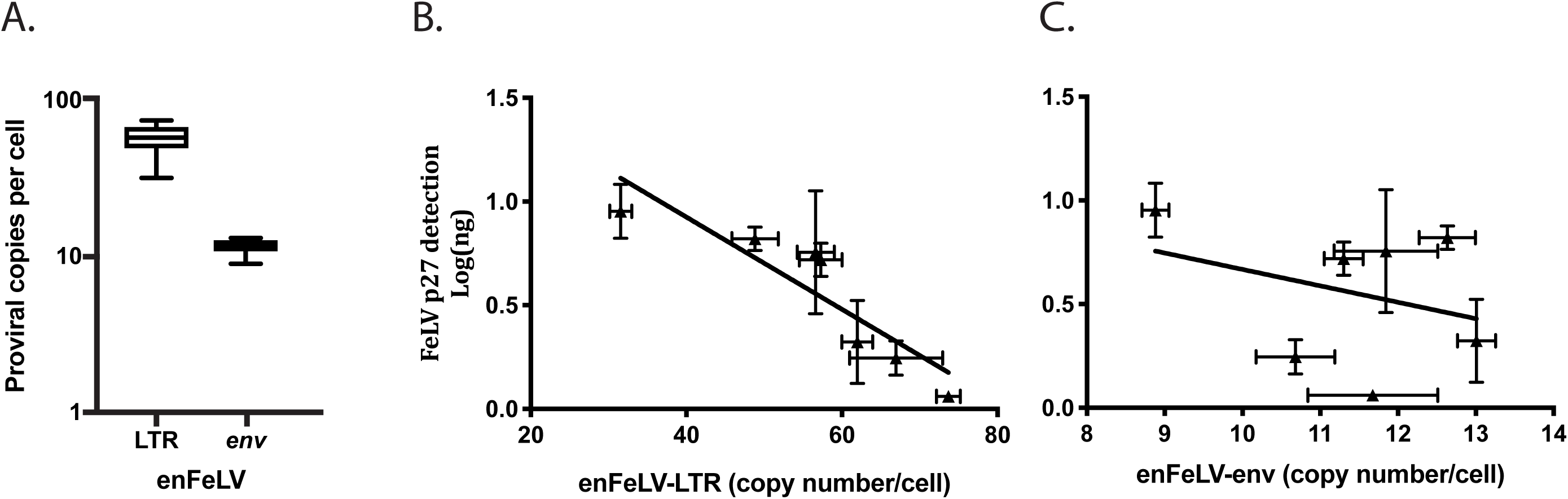
**A) enFeLV proviral loads vary between individual cats.** Domestic cat cells (n=7) display variation between enFeLV-LTRs and *env*. In individual cats, LTR copy number was greater than *env* copy number. enFeLV-LTR ranged between 9-13 copies per cell. enFeLV-env was more variable ranging between 32-74 copies per cell. **B) Domestic cat enFeLV-LTR copy number was negatively correlated with FeLV antigen production** (Pearson’s coefficient= −0.8943; p= 0.0066). **C) Domestic cat enFeLV-*env* copy number does not correlate with FeLV antigen production** (Pearson’s coefficient= −0.1071; p=0.8397), suggesting a direct role of enLTR in suppression of FeLV replication.

Variation in domestic cat fibroblasts enFeLV-LTR copy number correlated to FeLV antigen loads (day 7, Pearson’s correlation coefficient=−0.894; p<0.05; Figure 2B), whereas variation in enFeLV-*env* did not (day 7, Pearson’s correlation coefficient=−0.107, p=0.840; Figure 2C). enFeLV-exFeLV correlations were calculated at day 7 since this is the timepoint that cells reached complete confluency and the rate at which antigen was produced waned after this timepoint. Linear regression analysis of FeLV proviral load against antigen load showed that only 44% of the variation in antigen production could be explained by proviral load (R^2^ = 0.438; Figure 3).

**Figure 3.**
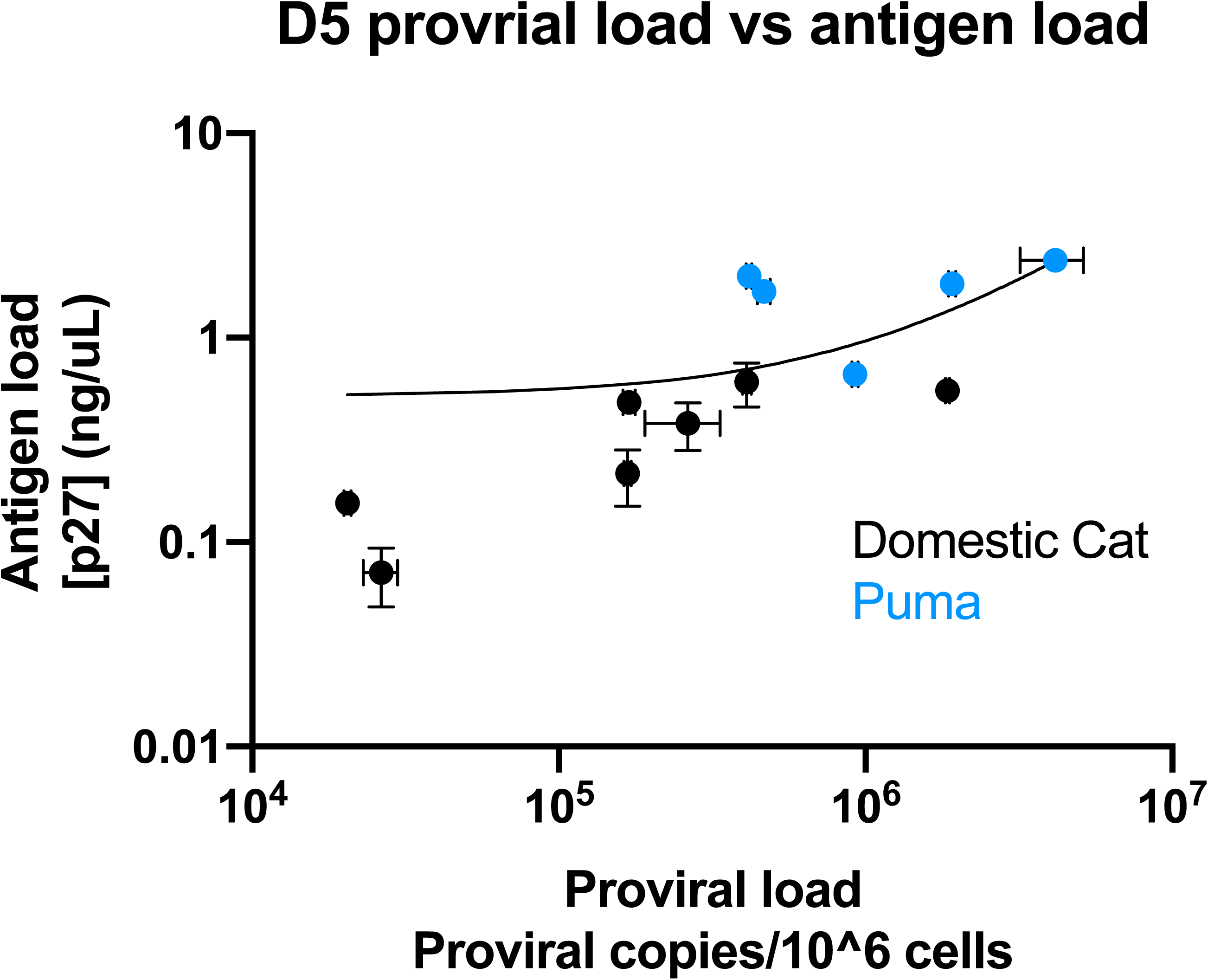
Proviral load does not highly correlate with viral antigen load. Linear regression analysis shows only 44% of variation in antigen production can be explained by proviral load following experimental FeLV infection of domestic cat (black) and puma (blue) cells. This indicates that the amount of viral antigen (a proxy for viral replication) is not governed completely by the degree of infection as measured by proviral load. Horizontal and vertical error bars indicate standard error for proviral load and viral antigen load, respectively.

### FeLV proviral and antigen loads during natural infection

FeLV loads in bone marrow, spleen, thymus and peripheral lymph nodes were assessed in 6 experimentally infected cats and 11 naturally infected pumas (not all tissues were available for each animal, see Table 1). Both tissue proviral load and tissue antigen load failed the Kolmogorov-Smirnov test for log-normality, indicating a non-normal distribution. FeLV proviral load was greater in domestic cats compared to panther by a median difference of 1.07 by Mann-Whitney test of log-transformed copy numbers per cell (U=122, p=0.0001; Figure 4A). Despite lower mean proviral load in Florida panther tissues, p27 capsid antigen in tissues tended to be higher (median value 3.41 vs. 2.87, Mann-Whitney test, U=250, p=0.127; Figure 4B). There was no difference in antigen or proviral load among tissues, with the exception of bone marrow antigen.

**Table 1.**
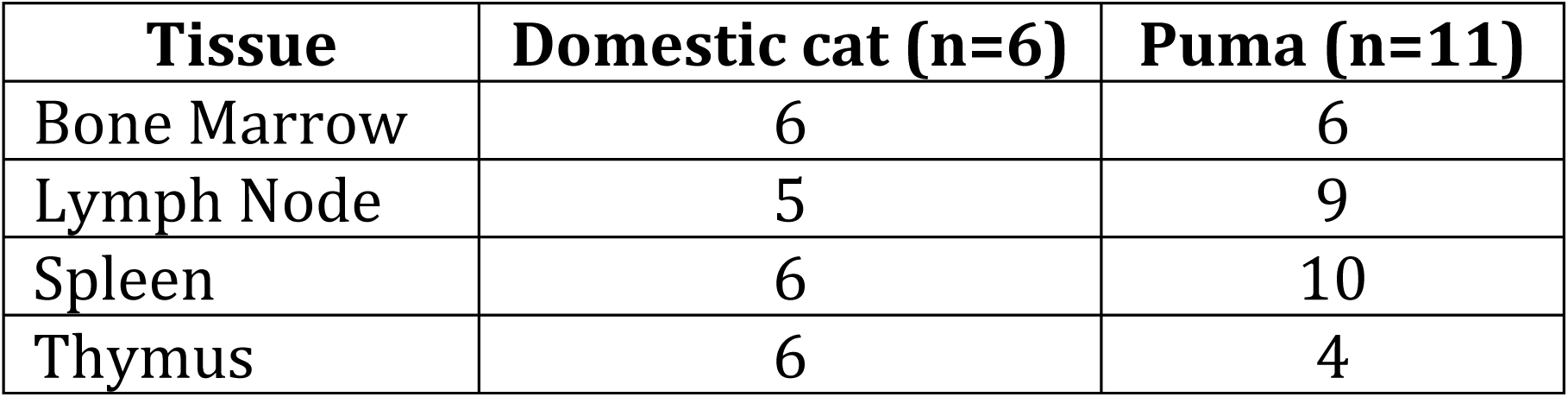
Tissues from FeLV-positive domestic cat and pumas used for qPCR and ELISA testing.

**Figure 4.**
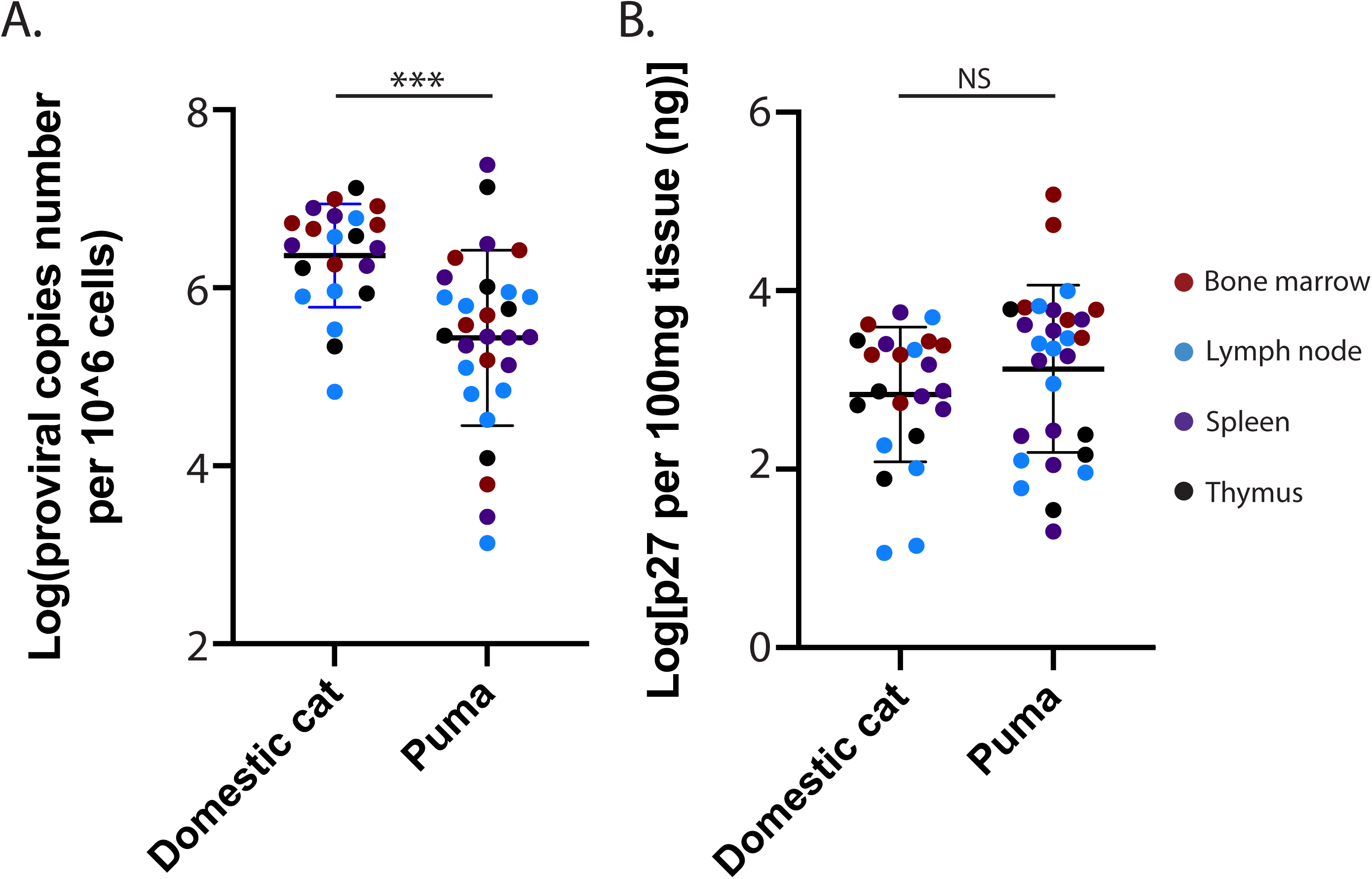
Lymphoid tissues from naturally infected FeLV-positive domestic cats and pumas have varying viral loads at end stage disease. A) Domestic cat lymphoid tissues displayed greater FeLV proviral load compared to puma tissues (Mann-Whitney test; ***p<0.0001). Puma tissues had a much wider range of viral loads which is potentially attributable to sample quality. B) Puma tissues trended to greater amounts of viral antigen (Mann-Whitney test; p=0.1495).

## Discussion

### Alterations in infectivity and virulence has been noted following cross species viral transmission (24)

In some cases, disease spillovers into novel species of the same family can result in dead-end hosts for the virus (Infectious hematopoietic necrosis virus (25); Feline immunodeficiency virus strain lru (26)). In other cases, disease spillover may result in active infections that maintain persistent transmission (Mycoplasmosis; (27); Feline foamy virus (28)). Further still, some cases result in adaptation of the virus in novel hosts leading to increased morbidity and mortality (HIV (29); Covid-19 (30)). Outcomes of diseases associated with spillover are dependent on the given specifics of host, environment and agent interactions (31). Multiple FeLV spillover events have been documented in free-ranging pumas with resultant significant morbidity (13, 32). The apparent virulence of FeLV in pumas and other nondomestic felids has led to speculation that this virus may have enhanced virulence in novel hosts (13, 14). This study was thus undertaken to evaluate the hypothesis that FeLV infection of nondomestic felids might be more competent in virus replication than the domestic cat reservoir host, examined specifically in the context of the presence of enFeLV.

### FeLV replicates more rapidly and productively in puma versus domestic cat fibroblasts

Experimental infections were conducted *in vitro* to establish differences in FeLV replication in puma and domestic cat cells in the absence of immunological and physiological parameters. In fibroblast cultures, FeLV infection resulted in higher proviral load in puma cells than in domestic cat cells (Figure 1A). Additionally, increased viral antigen was documented in infected puma cells, which is suggestive of increased viral production (Figure 1B). Therefore, at a cellular level, puma cells appear to be more competent at supporting FeLV infection and replication than fibroblasts of the primary host domestic cats. This is further supported by the fact that proviral load could only explain 43% of the variation in viral antigen production, indicating that proviral integration events alone are not a surrogate for virus replication. Other factors may be influencing the increased viral production in puma cells or restriction of viral replication in domestic cats

### enFeLV-LTR copy number is associated with resistance to FeLV infection and antigen production

Endogenous elements constitute a sizable component of an animal’s genome (2) and solo LTRs vastly outnumber full endogenous genomes/pseudogenomes (7). This occurs because two flanking LTRs allow for the intervening genes to be removed by homologous recombination, leaving behind just one copy of LTR it its place (33). As such, *env* copy number serves as a proxy for full enFeLV genomes, since loss of one gene would not occur frequently. In this sampling of domestic cat fibroblast enFeLV components, solo LTRs range from 32-74 copies per cell, while *env* ranges from 9-13 copies per cell, similar to previous observations (Figure 2A). This is consistent with previously reported measures of full-length enFeLV range of 6-26 copies per cell (23, 34, 35), versus 19-58 copies of enFeLV LTR per cell (23). LTR copy number variation, but not full enFeLV genomes, correlated with FeLV replication as evidenced by directly proportional antigen load; enFeLV-*env* showed no correlation to either FeLV replication (Figure 2B-C). While FeLV replication may intuitively seem to be correlated to proviral integration number, only 44% of FeLV antigen variation is explained by proviral load (Figure 3). Linear regression analysis therefore suggests that factors other than proviral integration number contributes to host susceptibility. This also corroborates observations made *in vivo* in a domestic cat colony naturally infected with FeLV (23), strongly suggesting that enFeLV LTR, versus other enFeLV elements, interacts with exogenous FeLV to mitigate infection.

CRFK proviral copy number and viral replication had a much smaller range than primary domestic cat and puma cell infections, with domestic cat variation exceeding individual puma range (Figure 1). This observation related to domestic cat cells can be explained by the constant enFeLV LTR copy number of this cell line (44 copies per cell). Greater variation in viral infection outcome in puma cells suggests that other host restriction factors modulate individual viral replication capacity in addition to enFeLV LTR.

One mechanism by which endogenous retroviruses may interact with exogenous retroviral infection includes the co-option of endogenous retrovirus long terminal repeats (LTRs), which harbor enhancer and promoter regions to stimulate host gene transcription. Endogenous retrovirus genome structures consist of protein encoding genes (*gag, pol*, and *env* are the minimum requirements for infectious retroviruses) capped on both ends by LTRs oriented in the same direction (4), since the process of recombination and retrotransposition, particularly early on in the endogenization process, can lead to the production and accumulation of additional LTRs scattered across the host genome (33). Retrotranspostion is important in allowing the virus to integrate its genome into other loci of the host genome (3) whereas the subsequent homologous recombination may leave solo-LTRs in potentially advantageous loci (33). LTRs may activate anti-viral or innate immune genes in close proximity to LTR insertion sites, result in protection against exogenous retroviruses. This indirect ERV modulation has been documented in other systems and can be mediated through *cis-*activation (promoter) or *trans-*activation (enhancer) of host genes (36). In the case of murine leukemia virus (MuLV), endogenous MuLV-LTRs have been co-opted to transcribe host antiviral genes including, but not limited to APOBEC3, a potent host retroviral restriction factor (37).

Alternatively, it is possible that the enFeLV genetic elements may directly interfere with exFeLV infection by encoding for small interfering RNAs or PIWI interacting RNAs that activate host DICER complexes to specifically target FeLV transcripts (38, 39). Our results are more suggestive of direct interference mechanisms due to the linearity of the FeLV restriction afforded by enFeLV-LTRs. It is unlikely that all LTR integration sites are influencing the transcription of host genetic factors and would have an effect in a dose-dependent manner.

### FeLV reaches high viral load in lymphoid tissues during natural infections

Naturally infected pumas with FeLV had lower proviral load than domestic cats, with the exception of two bone marrow samples, that achieved 2×10^7^ proviruses per 1×10^6^ cells (Figure 4A). Interestingly, a much wider range of proviral copy numbers were noted in pumas, and viral antigen loads in pumas equaled or exceeded that of domestic cats (Figure 4B). Field collections were performed opportunistically on Florida panthers when animals were either found deceased or hit by vehicle, often hours to days after death occurred. In contrast, FeLV positive shelter cats were euthanized prior to death, and tissues were collected rapidly following death. The timeliness of collection likely impacted quality of the sample prior to DNA extraction for puma samples. While normalization to feline CCR5 helped to address these issues for proviral copy number calculations, it is possible that viral antigen loads measured in puma tissues underestimate actual values.

Biological aspects of FeLV transmission that differ between domestic cats and pumas may impact subsequent infection kinetics. The initial spillover events of FeLV to pumas has been associated with predation of domestic cats (13, 40), whereas FeLV in domestic cats is believed to be transmitted in households through social interactions such as grooming, or via antagonstic interactions (11). Domestic cats interact socially, so behaviors like grooming may sustain infection in animals in close contact and infections may result from repeated exposures. Unlike domestic cats, pumas are much more solitary and interactions between pumas outside of mother-offspring groups are primarily believed to be antagonistic (41).

### No differences exist between cell culture parameters in relation to FeLV infection

Gammaretroviruses require the dissolution of the nucleus during mitosis in order to integrate into cells (42), and therefore dividing cells are more susceptible to FeLV infection and replication. We measured cell count as a proxy for rate of cell division, and percent cell mortality as a measure of viability. Neither measure differed between domestic cat and puma fibroblasts, though CrFK cells displayed greater cell count and cell mortality at confluency (Supplemental figure 1). Immortalized cell lines have accumulated multiple changes that fundamentally alter their morphology and physiologic behaviors, including decreases in contact inhibition (43). FeLV proviral integration and antigen production in CrFK infections had far less within- and between-run variation than primary fibroblasts, likely attributable to the clonal nature of CrFK versus wild type derived primary tissue cultures. Observation of cell culture parameters did not suggest differences in growth characteristics that explain the variant FeLV susceptibly of puma and domestic cat fibroblasts.

In this report, we present information that demonstrates that enFeLV-LTR confers protection against exFeLV infection *in vitro* through the limitation of FeLV replication. The exact mechanism by which these constituents act has yet to be determined, but leaves room for further investigation in the FeLV system as well as other endogenous-exogenous retroviral dyads. We hypothesize that FeLV restriction may manifest as direct interference through RNA silencing mechanisms, or by indirect enFeLV-LTR-mediated promotion of host anti-viral genes. FeLV provides an opportunity to directly interrogate the mechanisms that govern related exogenous-endogenous retroviral interactions in an outbred and diverse population.

## Methods

### Sample Acquisition

Full skin thickness biopsies from puma were collected by ear punch by the Colorado Parks and Wildlife under approved IACUC #16-2008. Abdominal skin samples were opportunistically collected from domestic cats during necropsies performed at the Colorado State University College of Veterinary Medicine. Primary fibroblasts were isolated as previously described (44) and cultured in 20% FBS-supplemented DMEM high glucose media and 1x antibiotic-antimycotic solution (Gibco, penicillin/streptomycin/fungizone). One puma culture was infected with feline foamy virus and was treated prophylactically with the anti-retroviral drug, AZT (100 ug/ml; Sigma) per manufacturer’s direction until in cultures where FFV CPE were detected. Cells were passaged two times for approximately 10 days in media without AZT washout prior to infection. Primary cultures were expanded for at most four passages before being frozen in (20% DMSO, 10% FBS, 70% serum free DMEM) using a freezing container (Nalgene) and stored at −80°C.

Bone marrow, thymus, spleen, and lymph node from naturally FeLV-infected domestic cats (n=6) and Florida panthers (*P. concolor coryi*, n=11) were collected and shipped to the Colorado State University Feline Retrovirus Research Laboratory for additional testing. Panther tissues were collected by the Florida Fish and Wildlife Conservation Commission from animals that had succumbed to FeLV following the introduction of the virus in two epizootics (13, 21). Domestic cat tissues were obtained from cats with terminal FeLV disease graciously provided by Animal Protective League (Springfield, IL). Puma and domestic cat lymphoid tissues were prepared for ELISA and qPCR assays as described below. DNA was extracted by bead-beater disruption and phenol-chloroform extraction using previously reported methods (13) to measure FeLV proviral load. Tissues were homogenized by bead-beater disruption (6.0 m/s for 60 seconds) in PBS with protease inhibitor (Pierce, Waltham, MA). Homogenates were diluted to 1% and 0.1% tissue for p27 capsid antigen quantification.

Mycoplasma-free Crandall Reese feline kidney cells (CRFK) were generously obtained by Dr. Martin Lochelt. Infections occurred when cells were between passage number 4 and 15.

### Virus titration

Crandell Reese feline kidney cells (CrFK) were plated at a density of 1,250 cells per 0.32cm^2^ in a flat-bottomed 96-well plate. FeLV-61E was obtained (gift of Edward Hoover and Candace Mathiason) and CrFK cells were infected in quintuplicate following a ten-fold dilution series. Cells were washed with sterile PBS, were given fresh media, and were incubated with 5% CO_2_ at 37°C for ten days. Titration was repeated three times. FeLV antigen ELISA detection, described below, was used to detect viral capsid antigen p27 in the supernatant. The quantity of virus necessary to infect 50% of tissue cultures (TCID_50)_ was calculated by previously published methods (45).

### enFeLV and exFeLV quantification by Real Time qPCR

LTR and *env* enFeLV copy number was quantified in domestic cat cells. *env* was used as a proxy for full-length endogenous FeLV, and LTR copy number detected both full length enFeLV as well as solo LTRs. Exogenous FeLV proviral DNA was measured by a third qPCR protocol targeting exFeLV specific LTRs, which vary from enFeLV (22). enFeLV-*env*, enFELV-LTR and exFeLV-LTR primers and probes were previously designed and reactions were performed as described (46) on a Biorad CFX96 thermocycler. In order to determine enFeLV and exFeLV proviral load, quantified FeLV was normalized against feline CCR5 (C-C chemokine receptor type 5; (47)) recognizing both domestic cat and puma CCR5 sequences. We used the delta cT method accounting for two CCR5 genes per cell (48). Custom DNA oligos were synthetically constructed with target regions of enFeLV LTR and env, exFeLV, and CCR5 on one DNA construct for quantification (gBlock, IDT; Supplemental figure 2).

Primer and probe sequences and qPCR thermocycling conditions are reported in Table 2 (22). exFeLV qPCR reactions contain 400 nM of both primers, 80 nM probe, iTaq Universal Probe Supermix (Bio-Rad), water, and DNA template. CCR5 exists as two copies per cell. Puma-specific CCR5 primers and probes adapted from Howard *et al*. were used to normalize FeLV copy numbers per 10^6^ cells (Table 2). The probe was labeled with FAM (6-carboxyfluorescein) reporter dye at the 5’ end, ZEN (Integrated DNA Technologies (iDT), Coralville, Iowa) internal quencher and IABkFQ (Iowa Black Fluorescein Quencher; iDT, Coralville, Iowa) at the 3’ end. CCR5 qPCR reactions contained 200nM forward primer, 500 nM reverse primer, and 200nM probe, iTaq Universal Probe Supermix (Bio-Rad), water, and DNA template. FeLV and CCR5 reactions were run simultaneously on the same plate on a Bio-rad CFX96 at 95°C for 3 minutes, followed by 40 cycles of 95°C for 5 seconds and 60°C for 15 seconds. The limit of detection for this assay is ≥10 copies per reaction. Standards for this assay were created as custom synthetic oligos (gBlocks, iDT) containing a relevant fragment of the exogenous FeLV and CCR5 genes (Supplementary Figure 2). Standard dilution and controls were run in duplicate and samples were run in triplicate.

**Table 2.**
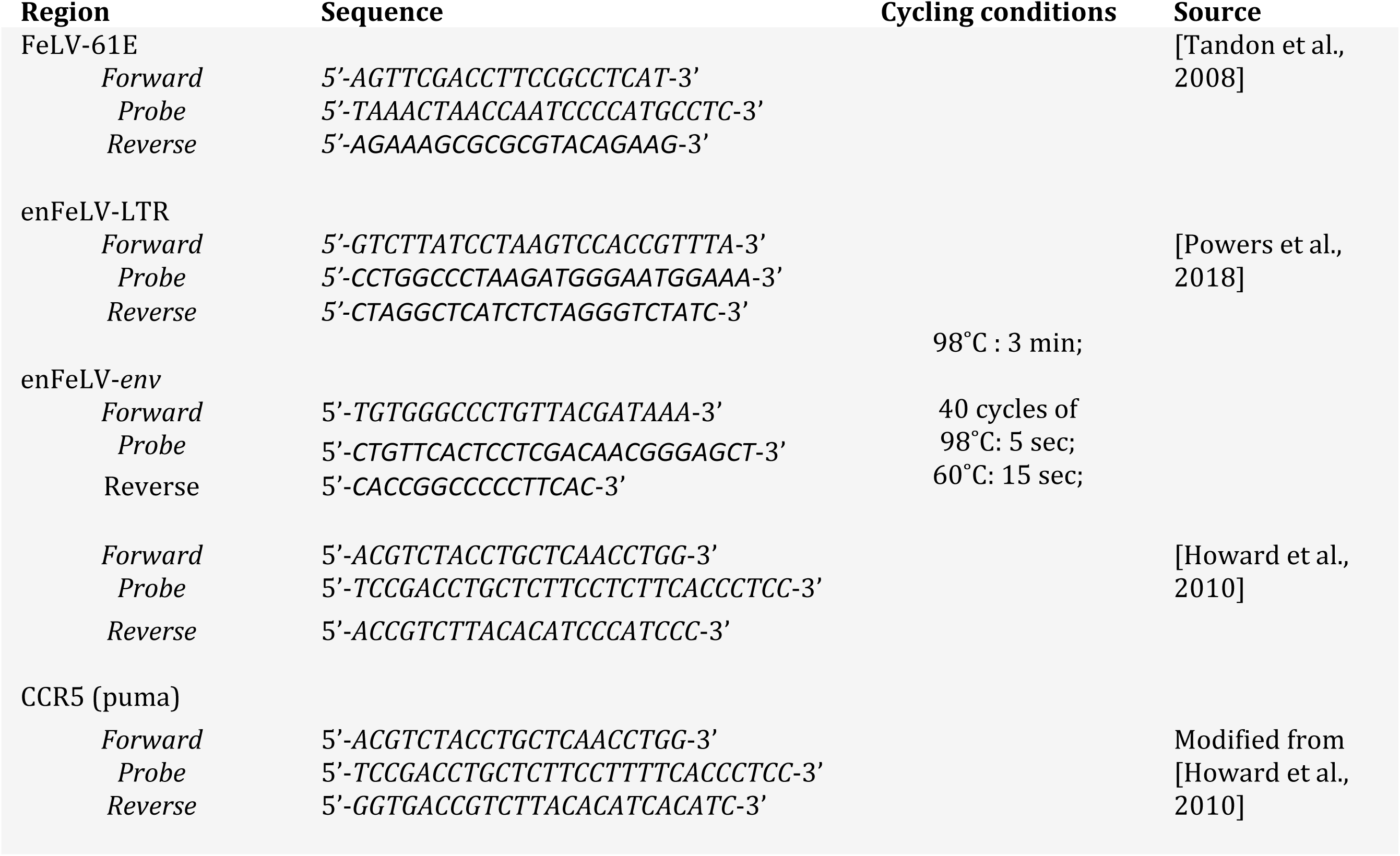
Primer sequences and cycling conditions used for qPCRs.

### Viral infection

Primary puma and domestic cat fibroblast cultures passaged fewer than five times were cultured in 20% FBS-supplemented DMEM high glucose media. Cells were plated at a density of 50,000 cells per 2 cm^2^ in a 24-well plate and infected with a multiplicity of infection (MOI) of 0.01 FeLV-61E in triplicate and cultured with 1.2mL media. 120 μL of supernatant was collected and stored at 80°C at days 0, 1, 3, 5, 7, and 10 for detection of p27 ELISA. At days 5 and 10, cells were harvested to determine cellular viability based on cell number and percent mortality by counting cells stained with trypan blue (Gibco) on a hemocytometer. One domestic cat cell culture, one puma cell culture, and one CrFK cell culture infection were terminated at day 7 due to equipment failure. One puma primary cell culture triplicate infection was repeated twice.

### ELISA

FeLV capsid antigen p27 was measured by sandwich ELISA. Costar Immulon 2HB plates were coated with 600ug CM1 capture antibody (Custom Monoclonal, Inc., US) in 100uL 0.1M Carbonate buffer (7.5 g/L Sodium Bicarbonate, 2.0 g/L Sodium Carbonate, pH ∼9.5) overnight at 4°C. Plates were blocked with 200uL 2% BSA in TEN buffer for two hours. One hundred μL of samples buffered with 50 μL ELISA diluent were incubated for two hours on a plate shaker. Six hundred micrograms of biotinylated secondary antibody (CM2-B; Custom Monoclonal, Inc., US) was incubated in each well, followed by 1:4000 dilution of HRP-conjugated streptavidin (ThermoFisherScientific, MA). Each step following sample incubation was followed with 5x wash with TEN buffer (0.05M TRIS Base, 0.001M EDTA, 0.15M NaCl, pH 7.2-7.4) with 0.1% Tween. All incubations were performed at room temperature. p27 antigen was detected indirectly following the addition of 3, 3’, 5, 5’ tetramethyl benzidine (TMB) substrate and peroxidase (Biolegend, San Diego, CA) at room temperature for 7.5 min before adding 2.5 N H_2_SO_4_ was quantified by Bioanalyzer at 450nm. Semi-purified FeLV p27 diluted in appropriate media (DMEM or RPMI) was used as a standard curve. Cutoff values for negative samples were three times the standard error over the average OD measured for control media samples.

## Acknowledgements

We would like to thank Dr. Lisa Wolfe, Mark Fisher, and Ivy Levan for collection and provision of puma tissue samples, Dr. Gary Mason, Dr. Jennifer Malmberg, Dr. Alex Byas, Dr. Laura Hoon-Hanks, Dr. Lauren Harris, and Dr. Devin von Stade for the collection of domestic cat full skin biopsies, and Esther Musselman and Mary Nehring for the collection of domestic cat blood.

This work was supported by NSF-EID award 1413925 and by the Office of the Director, National Institutes Of Health of the National Institutes of Health under Award Number T32OD012201 and F30OD023386. The content is solely the responsibility of the authors and does not necessarily represent the official views of the National Institutes of Health. The funders had no role in study design, data collection and interpretation, or the decision to submit the work for publication.

---

**Supplemental figure 1. Viable cell numbers and cell mortality increase over time and are similar in puma and domestic cat cultures.** A) Puma (blue) and domestic cat (black) fibroblast cell counts increased over time and reached 100% confluency by day 5. Initial seeding density was 5×10^4^ cells per 2cm^2^. Differences in domestic cat and puma fibroblast cell counts were not significant between day 5 and day 10. Minus signs denote uninfected cell cultures. B) Puma (blue) and domestic cat (black) fibroblast mortality increased over time following 100% confluency at day 5. Differences in domestic cat and puma fibroblast cell counts were not significant between day 5 and day 10. Minus signs denote uninfected cell cultures.

**Supplementary figure 2. Oligonucleotides synthesized for qPCR quantification.** Two gBlocks were synthesized commercially (Integrated DNA Technologies, Coralville, IA). Both concatenated gBlocks contained sequences for FeLV-61E (blue) and CCR5 (yellow). One contained enFeLV-LTR sequence, and the second contained enFeLV-LTR sequence. Bold sequences represent forward and reverse primer binding sites.

